# Carrier-frequency specific omission-related neural activity in ordered sound sequences are independent of omissions-predictability

**DOI:** 10.1101/2024.01.31.578194

**Authors:** Anne Hauswald, Kaja Rosa Benz, Thomas Hartmann, Gianpaolo Demarchi, Nathan Weisz

## Abstract

Regularities in our surroundings lead to predictions about upcoming events. Previous research has shown, that omitted sounds during otherwise regular tone sequences elicit frequency-specific neural activity related to the upcoming but omitted tone. We tested whether this neural response is depending on the unpredictability of the omission. Therefore we recorded MEG data while participants listened to ordered or random tone sequences with omissions occurring either ordered or randomly. Using multivariate pattern analysis shows that the frequency-specific neural pattern during omission within ordered tone sequences occurs independent of the regularity of the omissions suggesting that the auditory predictions based on sensory experiences are not immediately updated by violations of those expectations.

## Introduction

Even if people are not consciously aware of regularities in the environment, the brain does pick up on those and forms predictions accordingly. Such predictive processing can be seen in different modalities, e.g. touch (e.g. Dercksen et al., 2022; Yu et al., 2022), vision (Goujon et al., 2015), and hearing (Winkler et al., 2009). At a fundamental level, predictive processing enables us to form valid models of our environments (or updating them when necessary) as well as the consequences of our actions (Friston, 2010). An example par excellence is the learning of speech in early life, with studies showing the astounding statistical learning capacities of infants (Saffran, 2020). Abnormal predictive processing, especially an overweighting of predictions, may be a crucial mechanism that enables the development of phantom perception (Partyka et al., 2019; Sterzer et al., 2018). In many everyday settings, predictive processing especially in an anticipatory sense - i.e. relating to future events - enables more efficient processing, such as in iconic cocktail party settings (Schubert et al., 2023).

In auditory cognitive neuroscience, there has been a long and established tradition of experimental paradigms that probe predictive processes looking at event-evoked violations of regularity (so-called oddball-paradigms; Garrido et al., 2009; Schröger, 1998). Statistical regularity and by extension putative predictive auditory processing can be parametrically manipulated via so-called Markov sequences using e.g. pure tones (Barascud et al., 2016).

Applying multivariate pattern analysis techniques, including time- and condition generalization (King and Dehaene, 2014), we could recently show how increasing regularity of the pure tone sequence leads to increased feature specific neural activity (Demarchi et al., 2019). This was observed in particular prior to the onset of the anticipated sound (presented at 3 Hz) and following so-called omissions. The illustration of sound feature specific neural activity during “surprising” silent periods, goes beyond the demonstration of a general reaction of the brain, known as evoked or induced omission responses (Raij et al., 1997; Todorovic et al., 2011). For the latter, using a repetition suppression design, Todorovic and Lange (2012) showed the omission of a repetition lead to stronger responses when it was unexpected as compared to when it was more likely. Expectancy in this case was manipulated by overall frequency of the omission, either being frequent or rare depending on the condition. However, expectancy could also be manipulated via temporal predictability, keeping the overall frequency of omissions constant. Especially, how the temporal predictability of the occurrence of an omission influences the features specificity of neural responses as shown in our previous work (Demarchi et al., 2019), is unknown.

The goal of the present study is to address this open question. For this purpose, we presented regular and random tone sequences in which 10% omissions were embedded that were either strictly regular (i.e., every 10th tone) or appeared (pseudo-)randomly while recording MEG data. For ease of navigating through the manuscript, we will call the regularity of the tone sequence *frequency prediction* (F+) and the regularity of the omissions *temporal prediction* (T+), even though those terms are only rough approximations of the underlying processes. Our results show that regular sequences of tone frequencies are reliably linked to enhanced feature specific neural responses during omissions, irrespective of whether it was predictable or not. This suggests, predictions about upcoming tones are not reliably updated even though omissions violate those predictions (cf. stubborn predictions for visual cortex, Yon et al., 2023).

## Materials & Methods

### Participants

49 (23 female, 26 male) native German speakers participated in this study. Their age ranges from 18 to 37 (Mean = 23.32, SD = 3.66). Participants reported normal vision and hearing. Participation was voluntary and in line with the declaration of Helsinki and the statutes of the University of Salzburg All participants provided informed consent and were financially compensated for participation or via study credit. The study was approved by the ethical committee of the University of Salzburg.

### Stimuli and Procedure Procedure

Stimuli and procedure are similar to the ones in the study by Demarchi et al. (2019). Auditory stimuli were presented binaurally using MEG-compatible pneumatic in-ear headphones (SOUNDPixx, VPixx technologies, Canada) in sequences. These sequences were composed of four different pure tones, ranging from 200 to 2000 Hz, logarithmically spaced (200 Hz, 431 Hz, 928 Hz, 2000 Hz) each lasting 100 ms (5 ms linear fade in/out). Tones were presented at a rate of 3 Hz. Overall eight blocks were presented to participants, each containing 1600 stimuli. Each block was balanced with respect to the number of presentations per tone frequency (400 per tone). Within the block, 10% of the stimuli were omitted, thus yielding 400 omission trials (100 per omitted sound frequency). While within each block, the overall number of trials per sound frequency was set to be equal, blocks differed in the order of the tones and omissions. In more detail, there were four conditions, based on the order of tone sequence (ordered and random) and omission sequence (ordered and random). Random tone condition (see Fig. 1B) was characterized by equal transition probability from one sound to another, thereby preventing any possibility of accurately predicting an upcoming stimulus. In the ordered tone condition, presentation of one specific sound was followed with high (75%) probability by another specific sound thereby generating frequency-specific predictions. The probability for self-repetitions was set to 25% in both tone sequence conditions. The random omission condition had the omissions occurring randomly while in the ordered omission condition every tenth tone was omitted (balanced for the four different tones) thereby generating temporal predictions. Importantly, the absolute number of omissions was the same for both omission sequences. This resulted in four conditions in total (see Fig. 1A and C): frequency and temporal prediction (**F+T+**, when both tones and omissions occurred regularly), no frequency and no temporal prediction (**F-T-**, when both tones and omissions occurred randomly), frequency but no temporal prediction (**F+T-**, when tones occurred regularly but omissions randomly), and no frequency but temporal predictions (**F-T+**, when tones occurred randomly, but omissions regularly). During the stimulation a video of Cirque du Soleil (“Worlds Away”) was silently presented. The experiment was programmed in MATLAB 9.1 (The MathWorks, Natick, Massachusetts, U.S.A) using the open source Psychophysics Toolbox.

**Figure 1:**
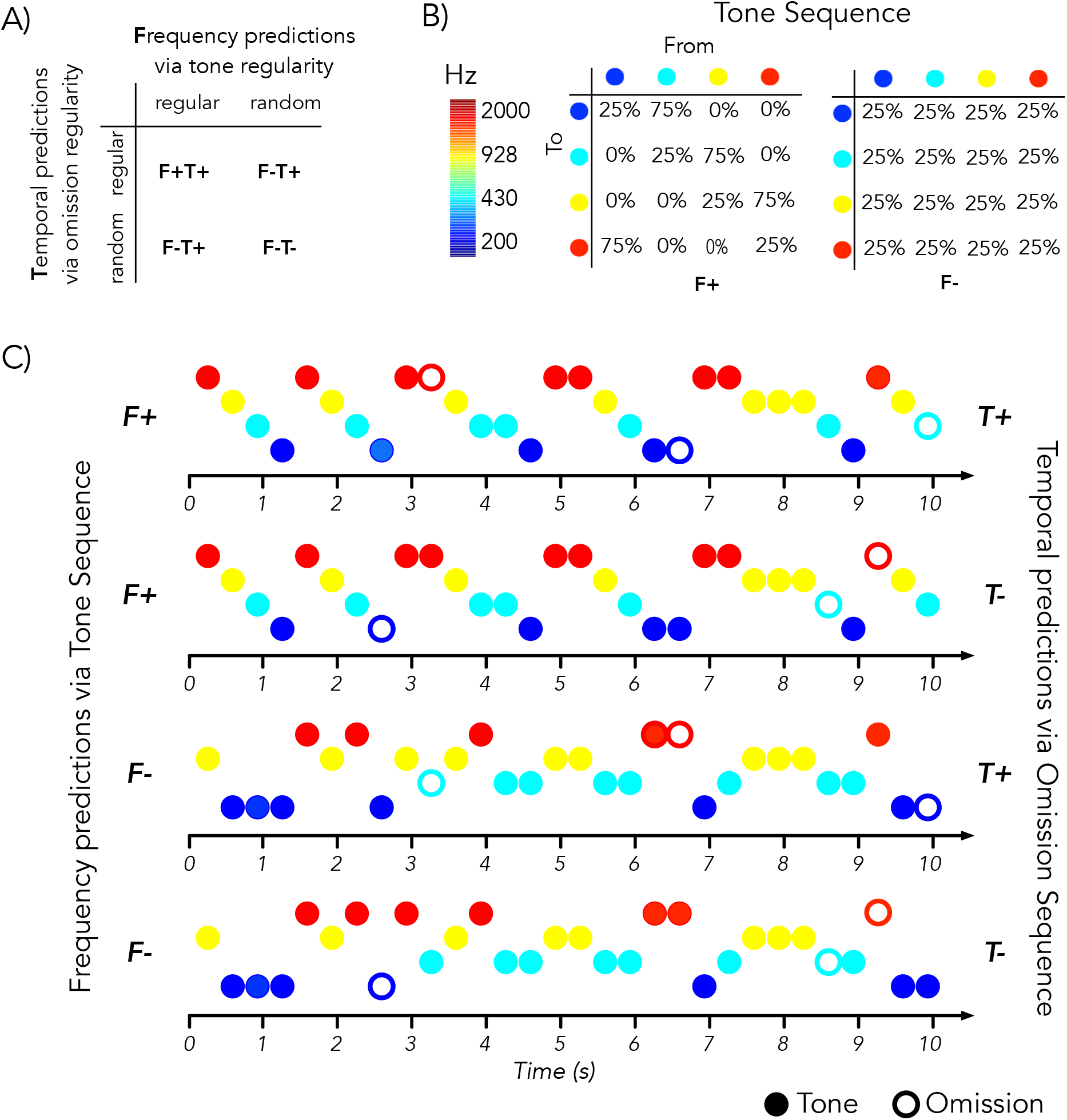
Overview of the experimental design. **A)** Illustration of the four conditions regarding tone and omission regularity **B)** Transition matrices of tones sequences in the different conditions **C)** Schematic examples of different sequences corresponding to the four conditions. The upper two sequences show regular tone sequences (F+) and the lower two random tone sequences (F-). The first and third tone sequence show sequences with regular omission (T+) and the second and last sequences random omission (T-). Omissions occurred in all conditions in 10% of the trials (absence of sound).

### Data acquisition and pre-processing

MEG data was recorded using a 306-channel Triux MEG system (Elekta-Neuromag Ltd., Helsinki, Finland) from 102 locations with 102 magnetometers and 204 planar gradiometers. In a magnetically shielded room (AK3B, Vakuumschmelze, Hanau, Germany). The MEG signal was recorded at a sampling rate of 1000 Hz and high-pass filtered at 0.1 Hz and low-pass filtered at 330 Hz. Prior to the experiment, the head shapes of each participant were digitized including fiducials (nasion, pre-auricular points) as well as around 300 points on the scalp using a Polhemus Fastrak Digitizer (Polhemus). We used a signal space separation algorithm as provided by the MEG manufacturer and implemented in the Maxfilter program (version 2.2.15) to remove external noise from the MEG signal (mainly 16.6 Hz from the nearby train and 50 Hz plus harmonics) and realign data to a common standard head position (across different blocks based on the measured head positions at the beginning of each block). Offline, data were analyzed using the Fieldtrip toolbox (Oostenveld et al., 2011). To reduce computational resources, only magnetometers were further analyzed. A 0.1 Hz high-pass filter was applied and the delay between trigger and stimulus onset due to the tubes delivering the acoustic signal was corrected. Data was cut into segments around the single tones and omissions (100 ms prestimulus and 300 ms poststimulus) and then resampled to 100 Hz.

### Decoding

All decoding was done using MVPA Light (Treder, 2020). Accuracy was calculated with a multiclass LDA classifier and the preprocessing settings of “demean” and “undersample”. Data was always trained on tones from the condition with random tone sequence and random occurrence of omissions (F-T-, non-overlapping trials with testing data).

For a first analysis of omission cross-decoding, testing data was chosen as trials corresponding to all F+ and F-omissions independent of the regularity of the omissions. We then also calculated the cross-decoding of the conditions separately for regularity of omissions and tone sequence, resulting in four conditions (random tones-random omissions F-T-, random tones-ordered omission F-T+, ordered tones-random omissions F+T-, ordered tones-ordered omissions F+T+).

### ERFs during omissions

Preprocessing for the omissions ERFs included a 1 Hz high-pass and a 30 Hz low-pass filter. Omission ERFs were calculated by timelocking data around the omission onset (-100 to 300ms) and using a baseline from -50ms to 0ms. Data were resampled to 250 Hz.

### Statistics

Decoding of all F+ omissions was contrasted with decoding of all F-omissions using cluster-based permutation (1000 repetitions, alpha 0.05). To compare the influence of the regularity of the omissions on this effect (i.e. temporal prediction), we extracted the individual differences of decoding accuracy between F+ and F-separately for T+ and T-for the maximal statistical value of the contrast between F+ and F-. We analyzed those differences with t-tests and with Bayes Factor. This was also one for the individual differences underlying the whole cluster.

Similarly, ERFs of all F+ omissions were contrasted to those of F-omissions using cluster-based permutations (0-300 ms, 1000 repetitions, two-tailed, alpha 0.05).

## Results

### Carrier frequency of omitted tones can be decoded in ordered tone sequences

In a first step, we wanted to replicate the key findings of Demarchi et al. (Demarchi et al., 2019) showing enhanced decoding accuracy of carrier frequency during the omission period in ordered tone sequences compared to random tone sequences. Therefore we contrasted the cross decoding of all F+ omission with F-omissions and yielded an effect around 100 ms training time, that lasted the entire omission period (testing time) with a peak around 100 ms (fig. 2 B upper panel), suggesting an effect of frequency prediction. F+ omissions yielded higher frequency-specific decoding than F-omissions (see also grandaverages in fig. 2A). This is similar to Demarchi et al (2019), although our effect is more restricted regarding training time (centres around 100 ms compared to 100-300 ms)

**Figure 2:**
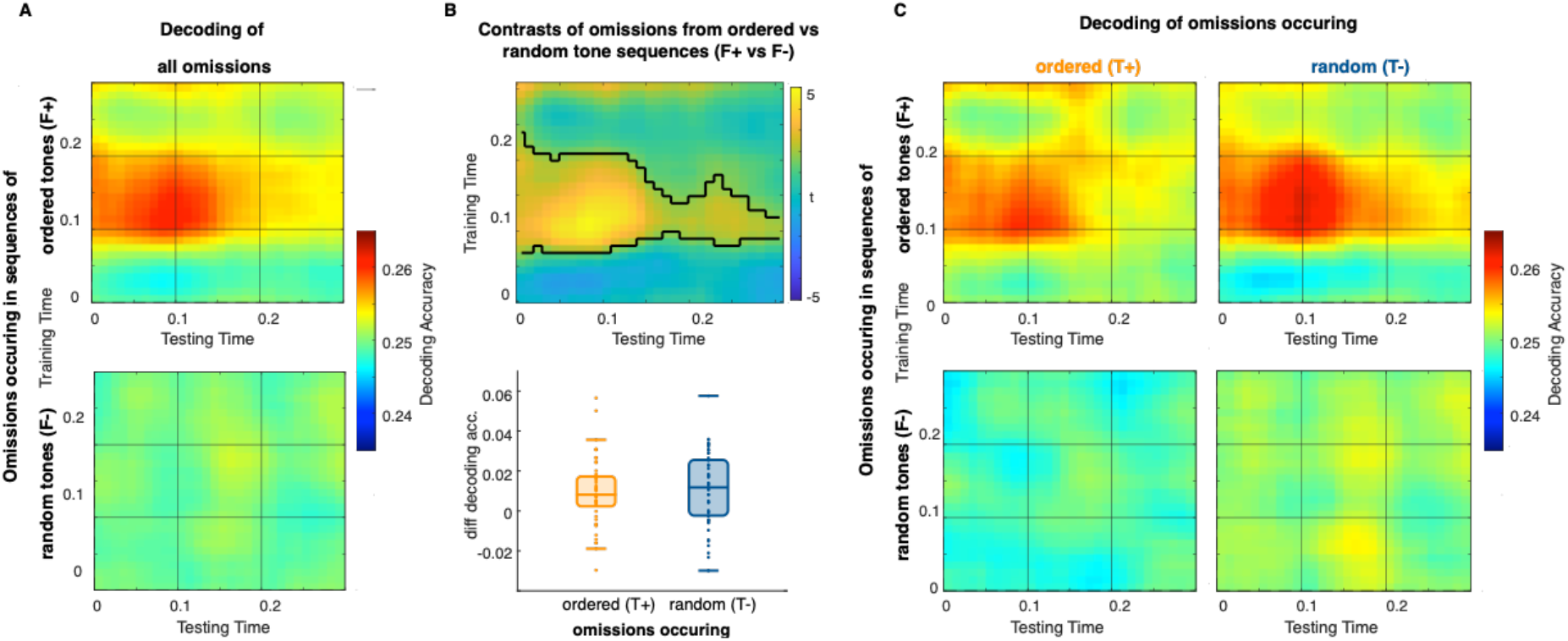
**A)** Grand averages of decoding accuracy of all omission trials from ordered tone sequences (F+, upper) and random tone sequences (F-, lower panel). **B)** Upper panel: Statistical contrast of decoding accuracy of all omission trials from ordered tone sequences (F+) and random tone sequences (F-). Lower panel: Boxplots of the difference between omissions within F+ and F-trials separately for omission occurring ordered (T+) or randomly (T-). **C)** Grand averages of decoding accuracy of all four conditions separately. Omission trials from ordered tone sequences (F+, upper) and random tone sequences (F-, lower panel), omissions occurring ordered (T+, left) and randomly (T-, right).

### Decoding of omitted tones is irrespective of expecting the omission

We then looked at the temporal predictions within those frequency predictions, i.e. looking at contrasts between omission within ordered and random tone sequences (F+ vs F-) separately for ordered (T+) versus random (T-) occurring omissions: no differences were found based on the regularity of the omissions using t-test (t(96)=-0.362, p-value=0.718). The bayes factor points to moderate evidence for the H0 (bayes factor= 0.1679, p-value=0.686 see boxplot in Fig. 2B and grandaverages in fig. 2C), i.e., that the temporal regularity of the omission occurrence does not influence sound frequency specific neural activity during the silent period. This result was stable also when using the whole statistical cluster to extract underlying individual values (t(96)=0.024, p-value=0.981, bayes factor= 0.213, p-value=0.981)

### No differences in omission ERFs in ordered and random tone sequences

Exploring the effects of frequency prediction on the evoked response similar to studies exploring MMN (Garrido et al., 2009) and repetition suppression (Todorovic and Lange, 2012), we contrasted the ERFs of all F+ omission with F-omissions. No significant differences (p-value >0.08) occurred, suggesting no effect of frequency prediction.

## Discussion

In this study we investigated how the temporal predictability of the occurrence of an omission influences the ability of the brain to form frequency-specific predictions of upcoming tones. For this purpose, we combined a paradigm of tone sequence regularity (markov sequence, e.g. Chait et al) with varying omission regularity and multivariate pattern analysis (Treder, 2020).

Omitting single tones within otherwise ordered tone sequences compared to random tone sequences offers the possibility to investigate the expectations about upcoming tones that are reflected in brain activity specific to the frequency of those upcoming tones. Using four levels of tone orderedness in a previous study, Demarchi et al. (Demarchi et al., 2019) showed that frequency-specific decoding of expected but omitted tones is dependent on the regularity of the tone sequence. The more ordered the sequence is, the stronger the frequency-specific decoding. In our study, we find a similar effect of increased decoding accuracy with a peak around 100 ms for omission periods of expected tones (F+) compared to not expected tones (F-). The statistical contrast shows that this increased decoding accuracy of F+ omissions lasts for the entire testing period of the omission (and around 100 ms training time) with a peak around 100 ms. Other studies have previously reported brain activity responding to a missing or deviant stimulus, e.g. in silent oddball paradigms (Busse and Woldorff, 2003; Karamürsel and Bullock, 2000) or studies presenting only one stimulus of previously paired associations (Bendixen et al., 2009; Stekelenburg and Vroomen, 2015). In line with Demarchi et al. (Demarchi et al., 2019), the present study shows, that the brain response to the missing stimulus, here missing tone, shows clear predictions of frequency-specific information.

The influence of the predictability of a missing tone itself however, is not clear as all omission occurred irregularly in the study by Demarchi et al. (2019) Therefore, in the present study, we manipulated the regularity of the occurrence of the omissions (temporal prediction) by having a random condition (T-) similar to Demarchi et al. (2019) and a regular condition (T+) where every tenth tone was omitted. Importantly, keeping the total number of omissions constant across conditions is different to e.g. Todorovic et al. (2012) using a repetition suppression design where the expectancy of omissions was modulated by often or rarely occurring omissions. In their study, the omission of a repetition led to stronger responses when it was unexpected as compared to when it was more likely.

Similarly, in Bekinschtein et al. (Bekinschtein et al., 2009) using the local-global paradigm global and local violations of regularity were modulated by frequency of occurrence.

To explore the influence of the temporal predictions of omissions on the frequency prediction of upcoming tones, we looked at the individual data underlying the maximal statistical values of the frequency prediction effect (fig. 2B) and calculated again the contrast of omissions from F+ and F-tone sequences separately for T+ and T-omissions. The temporal predictability of omissions has no influence on the frequency prediction as there is no change on the frequency-specific decoding due to the temporal prediction of the omission. This means, even if an omission is anticipated in an ordered tone sequence, the brain activity still captures the frequency-specific pattern of the omitted tone. This finding aligns with fMRI decoding results by Yon and colleagues (Yon et al., 2023), showing that perceivers hold an “undue reliance on old predictions” on visual outcomes without updating those based on unexpected and omitted visual outcomes. The idea of stubborn predictions explains the failing of the brain to update its predictions based on sensory evidence. As underlying mechanisms, computational limitations are discussed as well as estimation biases towards the predictive power of own actions. In the present study, another possible explanation for this undue reliance on established predictions might be that in an experimental paradigm, the tone-to-tone predictions in the ordered condition occur on a shorter time scale and therefore more often than the omissions (every 10th tone in the ordered condition) thereby differing in the number of learning trials.

Those stubborn predictions of the tone frequency are visible with decoding but not when looking at ERFs as there was already no reliable difference between responses during omissions within ordered tone sequences and those occurring within random tone sequences suggesting decoding to be more sensitive approach for identifying frequency-specific neuronal responses.

The absence of an effect for ERFs might be related to the length of the stimulus onset asynchrony (SOA). While we used a SOA of 300 ms, both animal studies (Lao-Rodríguez et al., 2023) and human MMN studies (Tervaniemi et al., 1994) have suggested that evoked responses might depend on fast tone presentations (<150 ms SOA).

Our study shows, that frequency-specific information is predicted by the brain during silent episodes within otherwise regular sequences and that this specificity is not dependent on the regularity of those silent periods. It further suggests by the absence of ERF related effects of frequency prediction and the presence of those effects for decoding that the decoding approach is more sensitive to illuminating effects of frequency specific prediction.

## Acknowledgements

Part of data acquisition was done as part of a course on experimental techniques.

## Author contributions

G.P. and N.W. designed the study. K.B. and T.H. acquired the data.

A.H. analyzed data. A.H. drafted manuscript. A.H. and N.W. edited the manuscript.

## Data accessibility

Processed MEG data will be made publicly available by time of publication

## Abbreviations

ERF: event related field
F: frequency prediction
fMRI: functional magnetic resonance imaging
MEG: magnetoencephalography
MMNI: Mismatch negativity
MVPA: multivariate pattern analysis
SD: standard deviation
SOA: stimulus onset asynchrony
T: temporal prediction

## References

Barascud, N., Pearce, M.T., Griffiths, T.D., Friston, K.J., Chait, M., 2016. Brain responses in humans reveal ideal observer-like sensitivity to complex acoustic patterns. Proc. Natl. Acad. Sci. 113, E616–E625. 10.1073/pnas.1508523113

Bekinschtein, T.A., Dehaene, S., Rohaut, B., Tadel, F., Cohen, L., Naccache, L., 2009. Neural signature of the conscious processing of auditory regularities. Proc. Natl. Acad. Sci. U. S. A. 106, 1672–1677. 10.1073/pnas.0809667106

Bendixen, A., Schröger, E., Winkler, I., 2009. I Heard That Coming: Event-Related Potential Evidence for Stimulus-Driven Prediction in the Auditory System. J. Neurosci. 29, 8447–8451. 10.1523/JNEUROSCI.1493-09.2009

Busse, L., Woldorff, M.G., 2003. The ERP omitted stimulus response to “no-stim” events and its implications for fast-rate event-related fMRI designs. NeuroImage 18, 856–864. 10.1016/s1053-8119(03)00012-0

Demarchi, G., Sanchez, G., Weisz, N., 2019. Automatic and feature-specific prediction-related neural activity in the human auditory system. Nat. Commun. 10, 3440. 10.1038/s41467-019-11440-1

Dercksen, T., Widmann, A., Noesselt, T., Wetzel, N., 2022. Somatosensory omissions reveal action-related predictive processing. 10.31234/osf.io/8aze5

Friston, K., 2010. The free-energy principle: a unified brain theory? Nat. Rev. Neurosci. 11, 127–138. 10.1038/nrn2787

Garrido, M.I., Kilner, J.M., Stephan, K.E., Friston, K.J., 2009. The mismatch negativity: A review of underlying mechanisms. Clin. Neurophysiol. 120, 453–463. 10.1016/j.clinph.2008.11.029

Goujon, A., Didierjean, A., Thorpe, S., 2015. Investigating implicit statistical learning mechanisms through contextual cueing. Trends Cogn. Sci. 19, 524–533. 10.1016/j.tics.2015.07.009

Karamürsel, S., Bullock, T.H., 2000. Human Auditory Fast and Slow Omitted Stimulus Potentials and Steady-State Responses. Int. J. Neurosci. 100, 1–20. 10.3109/00207450008999674

King, J.-R., Dehaene, S., 2014. Characterizing the dynamics of mental representations: the temporal generalization method. Trends Cogn. Sci. 18, 203–210. 10.1016/j.tics.2014.01.002

Lao-Rodríguez, A.B., Przewrocki, K., Pérez-González, D., Alishbayli, A., Yilmaz, E., Malmierca, M.S., Englitz, B., 2023. Neuronal responses to omitted tones in the auditory brain: A neuronal correlate for predictive coding. Sci. Adv. 9, eabq8657. 10.1126/sciadv.abq8657

Oostenveld, R., Fries, P., Maris, E., Schoffelen, J.-M., 2011. FieldTrip: Open Source Software for Advanced Analysis of MEG, EEG, and Invasive Electrophysiological Data. Comput. Intell. Neurosci. 2011, 1–9. 10.1155/2011/156869

Partyka, M., Demarchi, G., Roesch, S., Suess, N., Sedley, W., Schlee, W., Weisz, N., 2019. Phantom auditory perception (tinnitus) is characterised by stronger anticipatory auditory predictions (preprint). Neuroscience. 10.1101/869842

Raij, T., McEvoy, L., Mäkelä, J.P., Hari, R., 1997. Human auditory cortex is activated by omissions of auditory stimuli. Brain Res. 745, 134–143. 10.1016/s0006-8993(96)01140-7

Saffran, J.R., 2020. Statistical language learning in infancy. Child Dev. Perspect. 14, 49–54. 10.1111/cdep.12355

Schröger, E., 1998. Measurement and interpretation of the mismatch negativity. Behav. Res.Methods Instrum. Comput. 30, 131–145. 10.3758/BF03209423

Schubert, J., Schmidt, F., Gehmacher, Q., Bresgen, A., Weisz, N., 2023. Cortical speech tracking is related to individual prediction tendencies. Cereb. Cortex 33, 6608–6619. 10.1093/cercor/bhac528

Stekelenburg, J.J., Vroomen, J., 2015. Predictive coding of visual–auditory and motor-auditory events: An electrophysiological study. Brain Res. 1626, 88–96. 10.1016/j.brainres.2015.01.036

Sterzer, P., Adams, R.A., Fletcher, P., Frith, C., Lawrie, S.M., Muckli, L., Petrovic, P., Uhlhaas, P., Voss, M., Corlett, P.R., 2018. The Predictive Coding Account of Psychosis. Biol. Psychiatry 84, 634–643. 10.1016/j.biopsych.2018.05.015

Tervaniemi, M., Saarinen, J., Paavilainen, P., Danilova, N., Näätänen, R., 1994. Temporal integration of auditory information in sensory memory as reflected by the mismatch negativity. Biol. Psychol. 38, 157–167. 10.1016/0301-0511(94)90036-1

Todorovic, A. Lange, F.P. de, 2012. Repetition Suppression and Expectation Suppression Are Dissociable in Time in Early Auditory Evoked Fields. J. Neurosci. 32, 13389–13395. 10.1523/JNEUROSCI.2227-12.2012

Todorovic, A., van Ede, F., Maris, E., de Lange, F.P., 2011. Prior Expectation Mediates Neural Adaptation to Repeated Sounds in the Auditory Cortex: An MEG Study. J. Neurosci. 31, 9118–9123. 10.1523/JNEUROSCI.1425-11.2011

Treder, M.S., 2020. MVPA-Light: A Classification and Regression Toolbox for Multi-Dimensional Data. Front. Neurosci. 14.

Winkler, I., Denham, S.L., Nelken, I., 2009. Modeling the auditory scene: predictive regularity representations and perceptual objects. Trends Cogn. Sci. 13, 532–540. 10.1016/j.tics.2009.09.003

Yon, D., Thomas, E.R., Gilbert, S.J., de Lange, F.P., Kok, P., Press, C., 2023. Stubborn Predictions in Primary Visual Cortex. J. Cogn. Neurosci. 35, 1133–1143. 10.1162/jocn_a_01997

Yu, Y., Huber, L., Yang, J., Fukunaga, M., Chai, Y., Jangraw, D.C., Chen, G., Handwerker, D.A., Molfese, P.J., Ejima, Y., Sadato, N., Wu, J., Bandettini, P.A., 2022. Layer-specific activation in human primary somatosensory cortex during tactile temporal prediction error processing. NeuroImage 248, 118867. 10.1016/j.neuroimage.2021.118867

